# FITC Conjugated Polycaprolactone-Glycol-Chitosan Nanoparticles Containing The Longwave Emitting Fluorophore IR 820 For In-Vitro Tracking Of Hyperthermia-Induced Cell Death

**DOI:** 10.1101/273748

**Authors:** Piyush Kumar, Rohit Srivastava

**Affiliations:** Department of Biosciences and Bioengineering, Indian Institute of Technology-Bombay Powai, Mumbai 400076, India

## Abstract

Cancer theranostic agent IR 820 loses its bioimaging ability once therapy is initiated. At the end of therapy, it becomes difficult to track the cancer cells. To address this, FITC conjugated Polycaprolactone glycol chitosan IR 820 nanoparticles (FITC-PCLGC-IR NPs) has been synthesized for in vitro tracking of hyperthemia induced cell death. Two approaches, namely *ex situ* and *in situ* have been pursued FITC conjugation to PCLGC-IR NPs. Further comparisons were made to FITC encapsulated PCLGC-IR NPs in terms of biocompatibility, cellular uptake, photothermal mediated cell death and imaging with respect to laser treatment. We have shown that an 808 nm diode laser treatment did not affect the imaging ability of these NPs whereas cancer. Time scanned fluorescence shows the excellent photostability of this formulation for a maximum of 5 min. The detailed studies of these approaches summarize that FITC conjugation to PCLGC-IR nanoparticles is an effective nano-theranostic solution for image-guided photothermal therapy.

## 1. Introduction

Most commonly used the fluorescent theranostic agent in cancer theranostic is Indocyanine green (ICG)[3]. However, its poor aqueous solubility, low quantum yield in nanoformulation, faster degradation and lower noise to signal ratios hampers its use for future generation therapy. IR 820 has better noise to signal ratio, high photostability, and longer stability inside the physiological system [4]. Compared to metal based theranostics such as gold, these NIR dyes are biodegradable [5], consequently they have become an ideal choice for cancer theranostic. However, the limitation of these theranostic dyes, are their ineffectiveness to act as an imaging agent after laser irradiation.

To acquire the imaging ability, several fluorescent dyes have been used either in an encapsulated or a conjugated form in respective nanoformulations [6][7][8][9]. However, the main challenge in these conjugations is the pH dependent fluorescence of these dyes. The demand for pH dependent bioimaging has gained a lot of attention in recent past due to the different environment of normal and cancer cells. The cancer cells have acidic environment while normal cells have slightly basic environment pH 7 to 7.6. An FITC dye has emerged as an excellent choice owing to its excellent pH dependence nature. FITC fluorescence has been shown to increase in pH range 5.0 through 8 [10]. Owing to this property, FITC has been conjugated to gold, magnetic and silica nanoparticles for imaging [11][12][13][14]. However, their biodegradation, agglomeration, and clearance through the body are still debated. FITC has also been conjugated to polymers and polymeric particles containing ICG for bimodal imaging [15][16][17]However, their role in imaging during and after the photothermal therapy i.e. laser irradiation is not yet fully understood.

We have previously shown the significance of IR 820 and glycol chitosan (GC) in nanoformulation for effective cancer therapy [18][19]. IR 820 stabilized PCLGC nanoparticles retain its property as a chemotherapeutic delivery agent even after laser treatment while a GC enhanced the cell death by acting as an immunostimulatory molecule. In the current work, we have performed in-depth analysis of FITC conjugation/ adsorption to the PCLGC-IR nanoformulation and its role in bioimaging and photothermal therapy with respect to 808 nm laser irradiation.

## Experimental

### Materials

Polycaprolactone (PCL), poly (vinyl alcohol) 13-23 kDa, glycol chitosan (GC), IR 820 dye, Hoechst 33342, propidium iodide (PI) and fluorescein isothiocyanate isomer 1 (FITC) were purchased from Sigma-Aldrich. Acetone, dichloromethane, and methanol were purchased from Merck India Pvt. Ltd. Cell culture components such as DMEM, RPMI 1640, antibiotic-antimycotic solution, Phosphate buffered saline (PBS) and trypsin–EDTA solutions were purchased from Himedia Pvt Ltd Mumbai India. All the chemical and reagents used were of analytical grade. MDA-MB-231 breast cancer cells were purchased from National Centre for Cell Sciences (NCCS) Pune, India.

## Methods

### Formulation of FITC PCLGC-IR 820

#### Ex situ FITC Conjugation

FITC has been conjugated on prepared nanoparticles similar to the protocol described previously [20]. Briefly, 1 mg of FITC isomer 1 has been dissolved in 0.1 M Phosphate Buffer Saline, or carbonate buffer was slowly added on prepared PCLGC-IR 820 nanoparticles reported previously18 while stirring at low rpm in dark for overnight (1:10 ratio). Thereafter, FITC adsorbed PCLGC-IR nanoparticles were centrifuged and washed at 1000 rpm for 20 min to remove unbound FITC. The nanoparticles were kept at 4°C under dark condition.

#### In situ FITC Conjugation

In an *in situ* conjugation, FITC was conjugated to GC in carbonate buffer [21]. An FITC-GC conjugate was dialyzed to remove any unbound FITC.. FITC-GC was added to the aqueous phase containing poly (vinyl alcohol) (with a molecular mass of 13 to 23 kD). PCL and IR 820 were dissolved in a 2:1 (v/v) mixture of dichloromethane and methanol. The organic phase was slowly added to the aqueous phase containing PVA and an FITC-GC conjugate to form an emulsion (organic to aqueous phase ratio 1:3). The emulsion was further sonicated for 5 min and stirred at room temperature in the dark till the evaporation of the solvent. The nanoparticles were centrifuged at 14000 rpm for 20 min, washed thrice with Milli-Q water and were stored at 4°C under dark condition.

#### FITC Encapsulation

FITC was encapsulated into PCLGC-IR NPs as per standard protocol described previously [18]. Briefly, FITC isomer 1 and IR 820 have been added in an organic phase containing Dichloromethane and Methanol. The organic phase was slowly added in the aqueous phase containing GC solution. The emulsion was sonicated for 5 min and stirred at room temperature in the dark till evaporation of the solvent. FITC conjugated IR820 nanoparticles are obtained by washing the nanoparticles with water thrice times at 14000 rpm for 20 min and the nanoparticles obtained were preserved at 4 °C under dark condition.

### Characterization of FITC PCLGC-IR

The NPs were characterized by Dynamic light scattering (DLS) and Zeta analyzer (Brookhaven Instruments Corporation, USA) and Scanning Electron Microscope (SEM; JSM 7600 F Jeol, Japan). The absorption spectra were recorded using UV-Vis spectrophotometer (Perkin Elmer USA). Briefly, 200 µL of nanoparticles of 1 mg/ mL concentration were subjected to wavelength scan from 400-1100 nm. To analyze FITC conjugation, fluorescence spectra of nanoparticles were recorded in Shimadzu fluorescence spectrophotometer (Japan) with λex 485 nm and λex 500.nm IR 820 nanoparticles in Milli-Q were taken as a control.

### Effect of Laser on FITC Conjugated Nanoparticles

Effect of the laser (808 nm for 5 min) was studied on FITC conjugated nanoparticles by UV-Vis spectroscopy and fluorescence spectroscopy. Briefly, 200 µL of nanoparticles of 1mg/ mL concentration were incubated at 37 °C prior to laser application. Two set of experiment (each n = 3) were performed to analyze the effect of absorbance and fluorescence separately. The laser was continuously applied to the sample for 5 min. At the end of laser treatment, samples were scanned for absorbance and fluorescence by UV-Vis and Fluorescence spectroscopy respectively.

### Photothermal Effect on NPs

Photothermal effect of nanoparticles was studied with 808nm infra-red laser (0.5 W). Briefly, 200 µL of 1mg/ mL suspension of nanoparticles was exposed to 808 nm laser in a 96-well plate in the water bath maintained at 37 °C. The temperature was monitored with Okaton digital thermometer. The rise in temperature was measured at the interval of 1 min up to 5 min (n = 3).

### Biocompatibility study

Mouse Fibroblastic cells (type NIH3T3) in a concentration of 1x10^4 cells/ml were seeded into a 96-well plate and incubated at 37°C in humidified 5% CO_2_ incubator for 24 h.. The medium was replaced with fresh complete media containing NPs from 100-1000 µg/ mL. The plate was further incubated for 24 h. After that, cell viability was determined by MTT assay (n = 3).

### Photothermal therapy

Breast cancer cells (type MDA MB 231) in a concentration of 1x10^4 cells/ml were seeded into a 96-well plate and incubated at 37°C in humidified 5% CO_2_ incubator for 24 h. Thereafter, cells were washed and incubated with fresh media with FBS (10%) containing 125 µg of NPs and incubated for 3 h. The photothermal effect on MDA-MB 231 cells was studied as per the protocol described earlier [18] (n = 3).

### Hoechst Propidium Iodide doubles staining

For live/ dead, apoptotic and necrotic cells visualization, Breast cancer cells (type MDA MB 231) in a concentration of 1x10^4 cells/ml were seeded on coverslip in 12 well plate and incubated at 37°C in humidified 5% CO_2_ incubator for 24 h. The remaining protocol was similar to photothermal effect as described earlier.[18]

### in vitro bioimaging

Breast cancer cells (type MDA MB 231) in a concentration of 1x10^4 cells/ml were seeded into a 96-well plate and incubated at 37°C in humidified 5% CO_2_ incubator for 24 h. After 24 h, cells were washed with PBS (pH 7.4) and incubated with media containing NPs for 3 h. The cells are washed thrice with PBS to remove any free NP and observed under Confocal laser scanning microscope (Olympus) at λex 485 nm and λex 520. nm (n = 3).

## Results and discussion

In our previous study, we have previously shown the degradation of IR 820 dye upon laser 808 nm irradiation [18]. Hence it could not be used as an imaging agent during and after laser treatment for effective cancer therapy which is the major drawback of most of the cancer theranostic agent. To retain the imaging ability of these NPs after laser irradiation, we have conjugated FITC in two ways: essentially via *ex situ* and *in situ* thereby comparing it with FITC encapsulated NPs for the first time

(1) In the first case, we prepared PCLGC-IR nanoparticles and followed by surface conjugation of FITC (*ex situ*).

(2) In the second case, the efficacy of conjugation was compared by first, conjugating FITC to glycol chitosan (GC) followed by nanoparticles fabrication by the solvent evaporation method *(in situ*).

We further prepared FITC encapsulated PCLGC-IR NPs (*en*) to compare among the all the three methods (Fig. 1). An FITC addition will provide an additional advantage over the nanoformulation wherein PCL being hydrophobic can be used encapsulation of drug and GC has been shown to increase the fluorescence [22], followed by IR 820 for photothermal ablation. The purpose of studying three different types of FITC incorporation was to compare the effectiveness and suitability of stable nanoformulation for effective post laser imaging. Hence, the NPs were further characterized by SEM for morphological, UV-Vis and fluorescence spectroscopy for stability studies.

**Figure 1:**
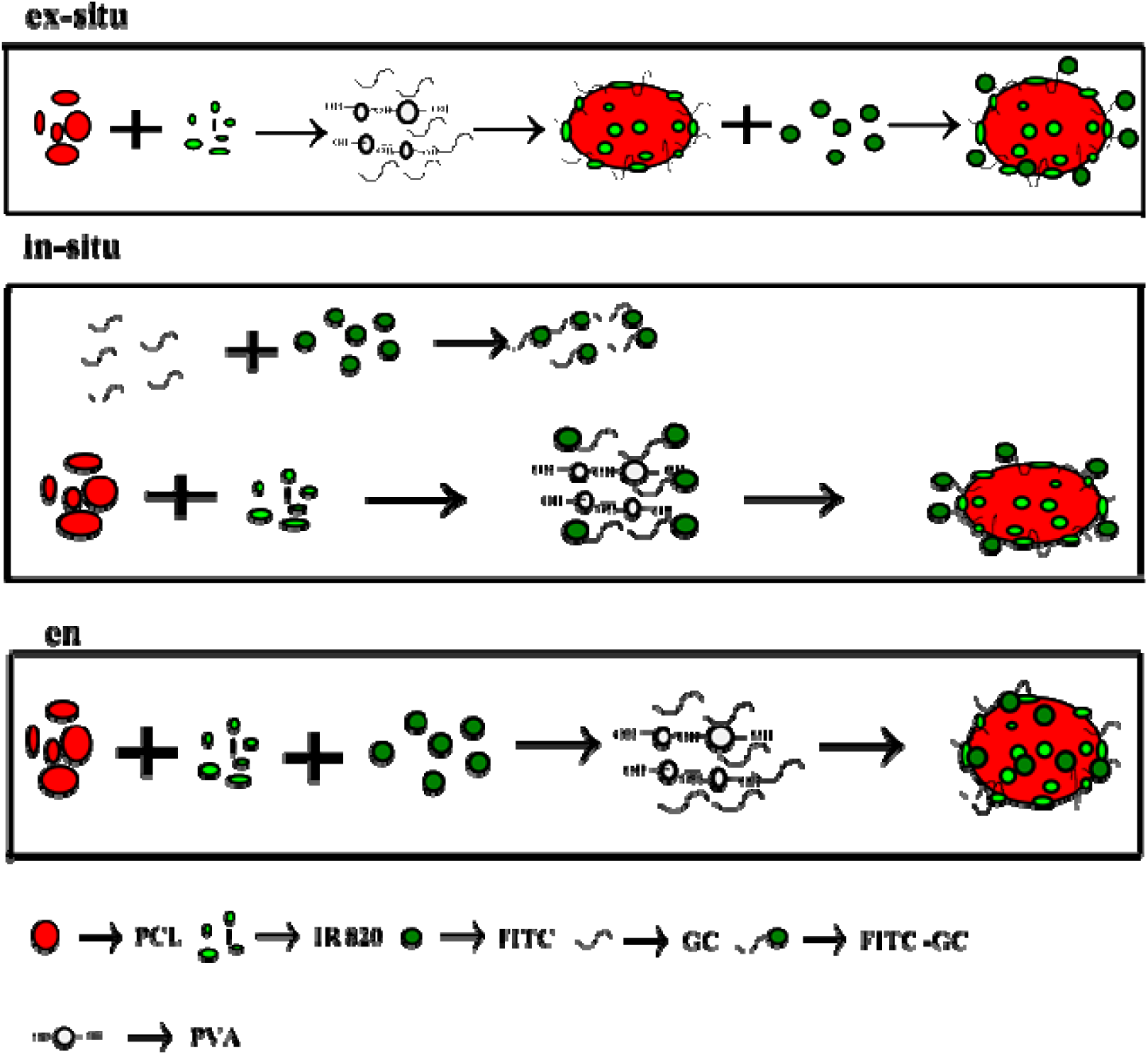
Schematic of FITC labelled PCLGC-IR nanoparticles.

The SEM image showed the size of both *ex situ* and *in situ* FITC NPs were around 150-260 nm while *en* NPs showed smaller size around 100-200 nm (Fig. 2). Both *in situ* and *en* NPs showed uniform monodisperse nanoparticles, while, *ex situ* FITC NPs tend to settle due to low zeta value. *Ex situ* NP has a zeta potential of around +2 mV whereas both *in situ* and en NPs showed zeta potential around +25 mV. High surface charge and uniform shape stabilized the *in situ* and *en* NPs in aqueous suspension. The positive charge even upon FITC incorporation can be used the attachment and monitoring of gene delivery.

**Figure 2:**
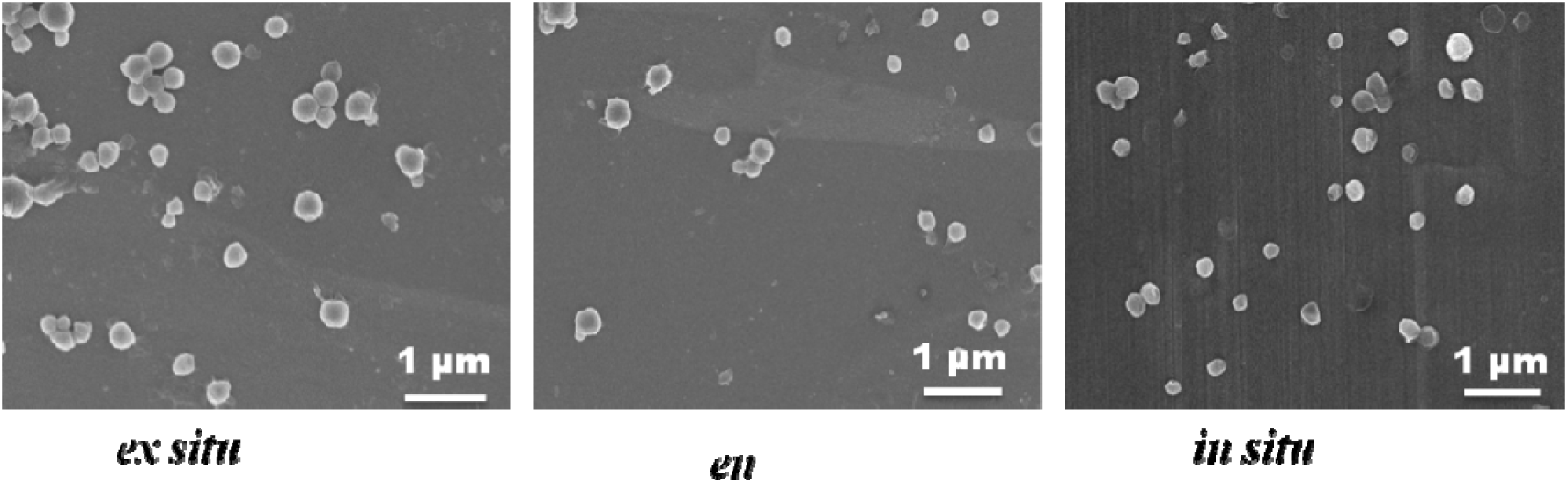
SEM images of FITC –PCLGC-IR NPs.

UV-Vis absorption spectroscopy was performed to analyze the FITC conjugation and its effect on IR 820 encapsulation (Fig. 3). PCLGC-IR has its characteristic peak around 876 nm and 750 nm. The position of the IR 820 peak did not change upon FITC conjugation/encapsulation. Both *in situ* and *ex situ* conjugation to the nanoparticles did not show any significant FITC absorption peak around 493 nm at a low concentration while *en* NPs showed the prominent peak around 493 nm inferring high encapsulation of FITC in *en* NPs. To further validate FITC conjugation, fluorescent measurement was performed using fluorescence spectroscopy. FITC fluorescence was much higher in *en* conjugation and *in situ* than that of *ex situ* PCLGC-IR.

**Figure 3:**
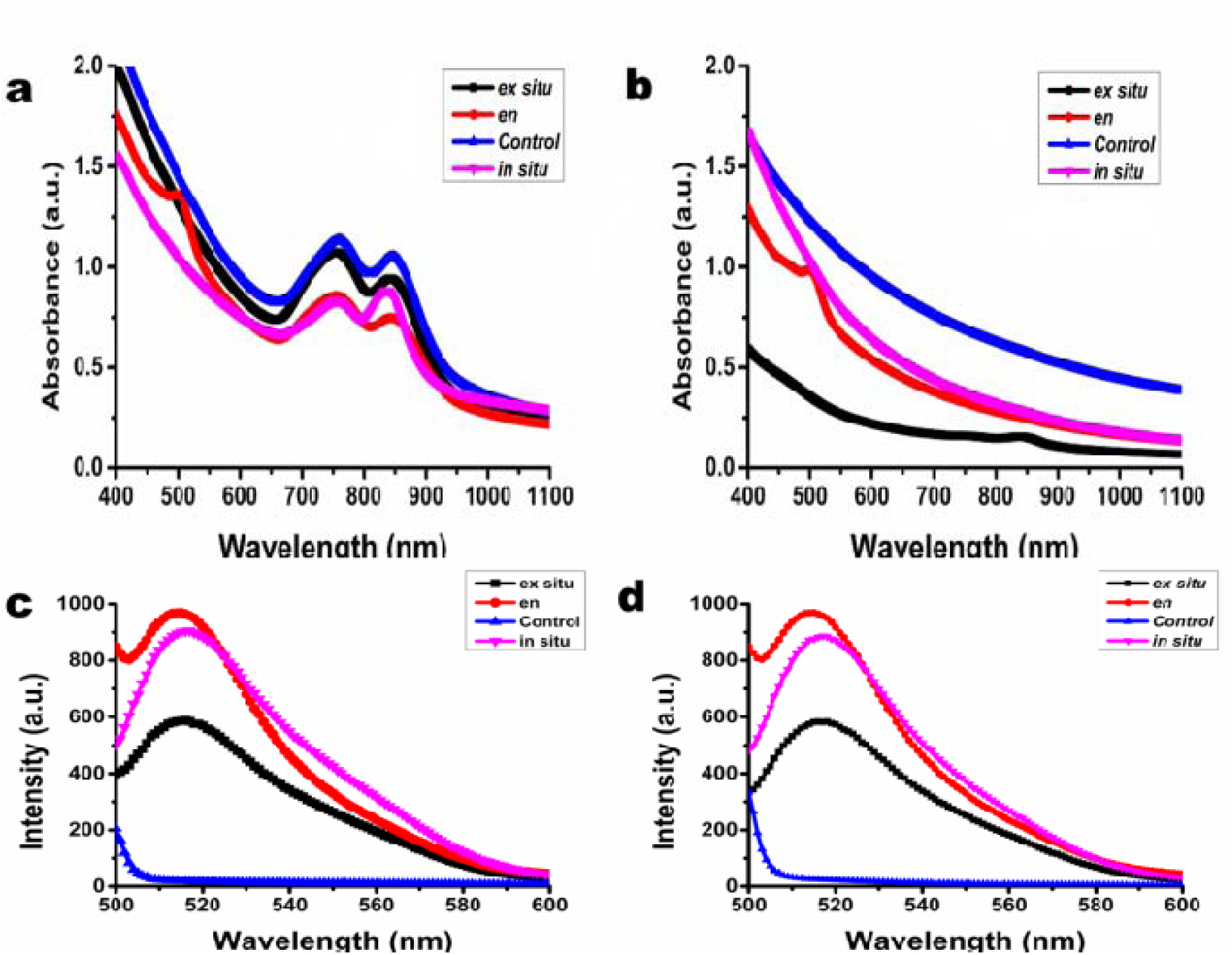
UV-Vis and fluorescence spectra of FITC -PCLGC-IR NPs (a) UV-Vis spectra of nanoformulations (b) UV-Vis spectra of 808 nm laser treated samples, (c) Fluorescence spectra of nanoformulation at excitation of 480 nm and emission at 500 nm and (d) Fluorescence spectra of laser treated samples at same excitation and emission wavelength. PCLGC-IR nanoparticles have been taken as a control.

### Effect of laser irradiation and stability of nanoformulations

Feifan Zhou et al have shown that FITC fluorescence diminished after laser irradiation [23]. To analyze the effect of laser on FITC conjugation, we further checked the UV-Vis absorbance of the laser treated samples which showed that IR820 peak was diminished in all cases (Fig. 3 b). However, there was no significant effect of FITC absorption in either of the conjugations/encapsulation. To rule out the the possibility of the effect of laser irradiation on FITC, fluorescence spectra were also recorded (Fig. 3c and 3d). Surprisingly, there was no change in fluorescence spectra of FITC in either of the cases of FITC labeling. This showed the superiority of our nanoformulation over the previously reported data. Also, time course fluorescence spectroscopy was performed for 5 min to analyze the FITC stability in all three nanoformulations (Fig. 4). All the three formulations were stable in given time period although *ex situ* showed some fluctuation and a decrease in intensity over the time as it tend to settle down. The particles tend to settle with time leading to random fluctuation in fluorescence scattering. The good stability of FITC in these NPs throws light on its application during photothermal therapy in the physiological system, wherein the green color of FITC labeled tissue can be observed over the red color of the blood for the optimal time.

**Figure 4:**
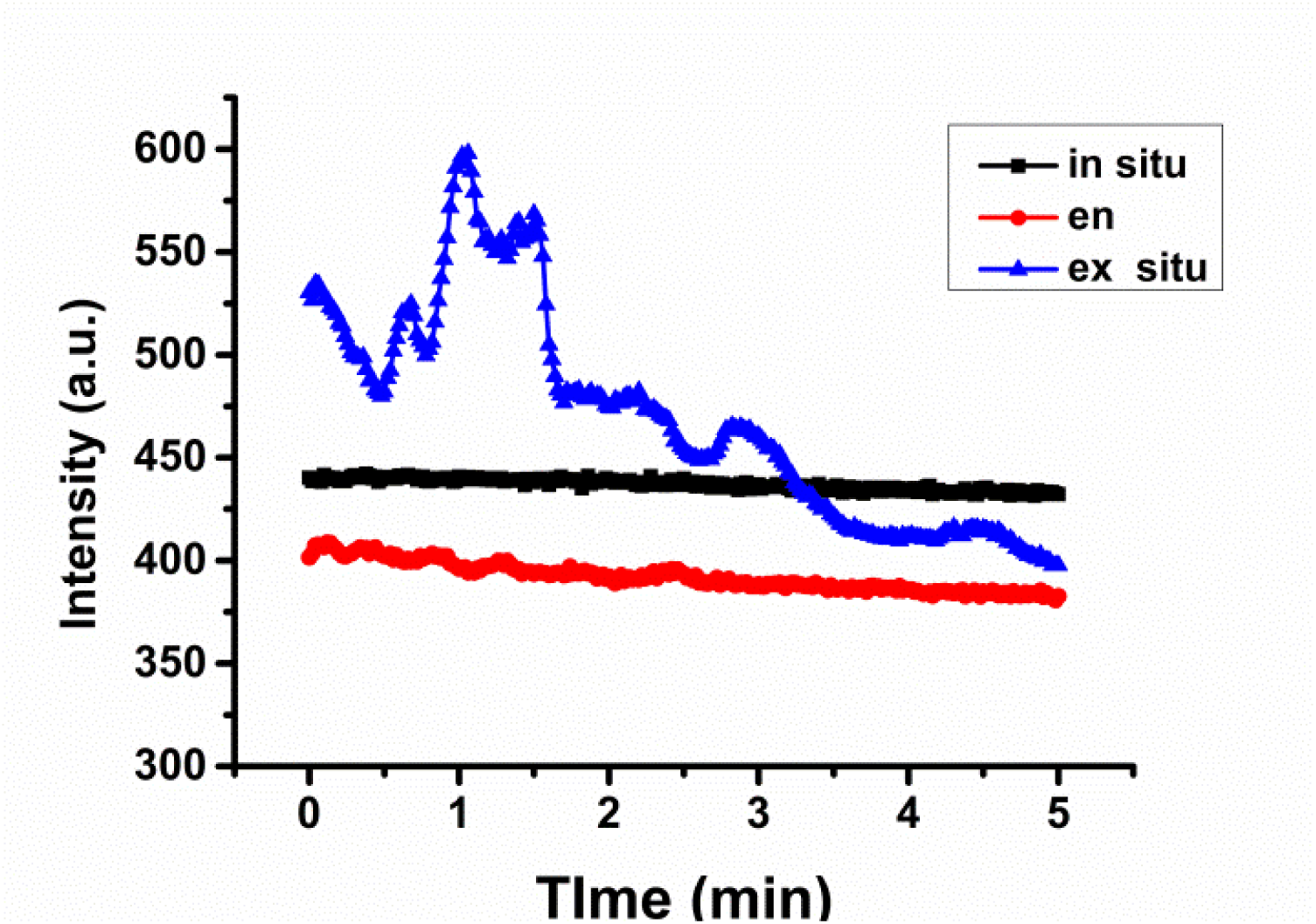
Time scan of FITC-PCLGC-IR NPs for 5 min.

### Photothermal transduction

We further analyzed the photothermal efficiency of FITC conjugated nanoparticles. The photothermal efficiency was higher in *ex situ* FITC conjugated to nanoparticles compared to both *in situ* and en NPs. The maximum temperature rise in *ex situ* conjugation was around 45°C while *in situ* and en showed around 43 °C and 42 °C, respectively (Fig. 5). These NPs retained the photothermal transduction property even after FITC integration. FITC protein conjugate has been shown to have poor stability at 37 °C and elevated temperature [24]. However, in our case, there was no difference in FITC fluorescence for 5 min as described earlier even the maximum temperature reached 45 °C. Another reason of FITC stability inside nanoformulation could be high zeta value that keeps the nanoparticles apart.

**Figure 5:**
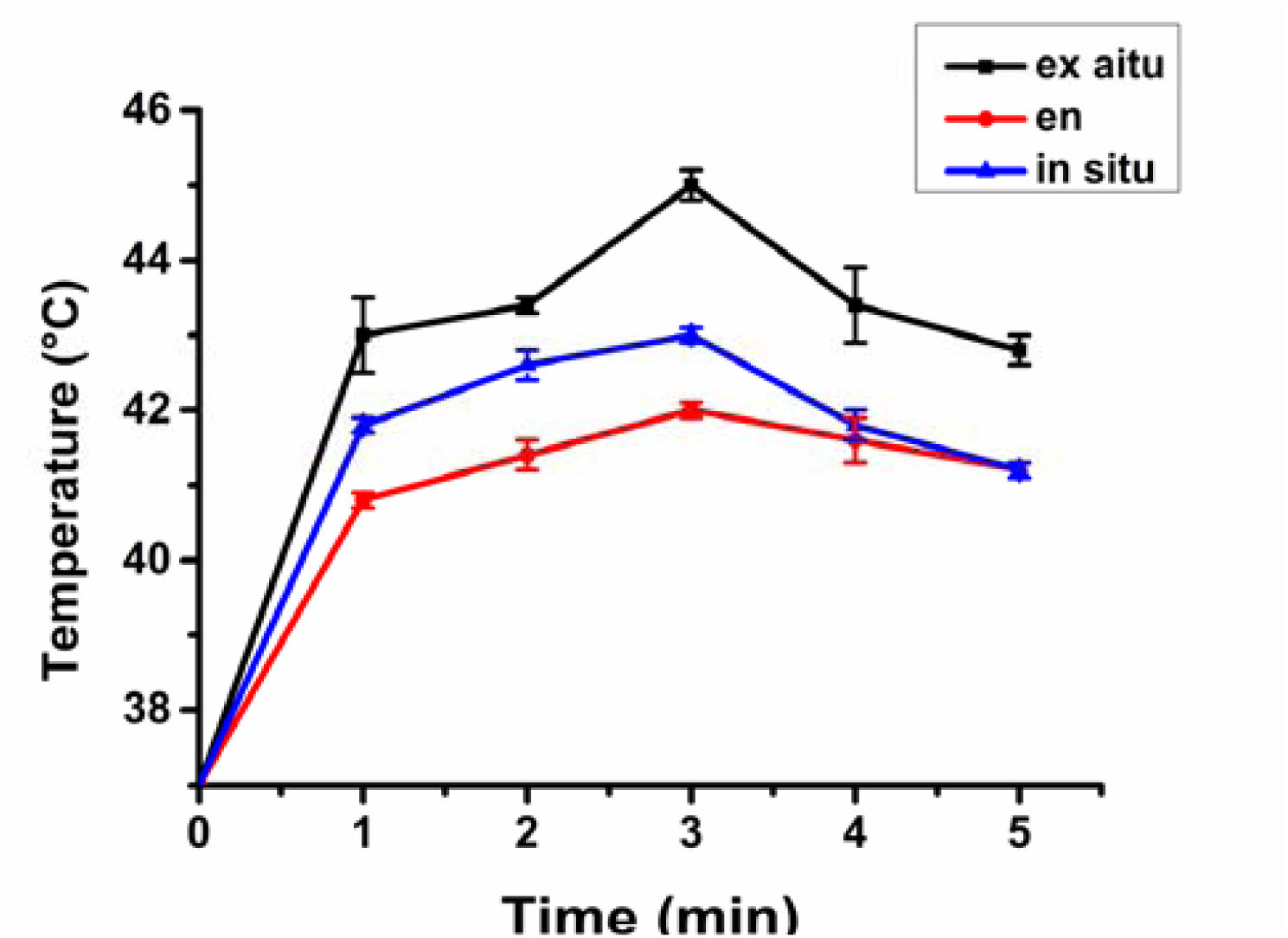
Photothermal transduction of FITC-PCLGC-IR NPs.

### In vitro cytotoxicity assessment

Cytotoxicity of the nanoformulation is a major concern in effective cancer treatment. The cytotoxic effect of FITC conjugated NPs was quantified by an MTT-based cytotoxicity assay on NIH3T3 cells (Fig. 6 a). All three were found biocompatible even at high concentration. We further analyzed the photothermal efficiency of these nanoparticles on MDA-MB-231 breast cancer cells (Fig. 6 b). An *ex situ* FITC conjugated to nanoparticles showed less photothermal efficiency in MDA-MB-231 cells compared to *in situ* conjugation and *en* NPs. To visualize the cell death, we further stained the cells with Hoechst 33342 and PI (Fig. 6 c). The HPI double staining showed cell death upon laser irradiated NPs treated cells in all the formulation. *In situ* conjugated NPs formation showed good cell death compared to *ex situ*.

**Figure 6:**
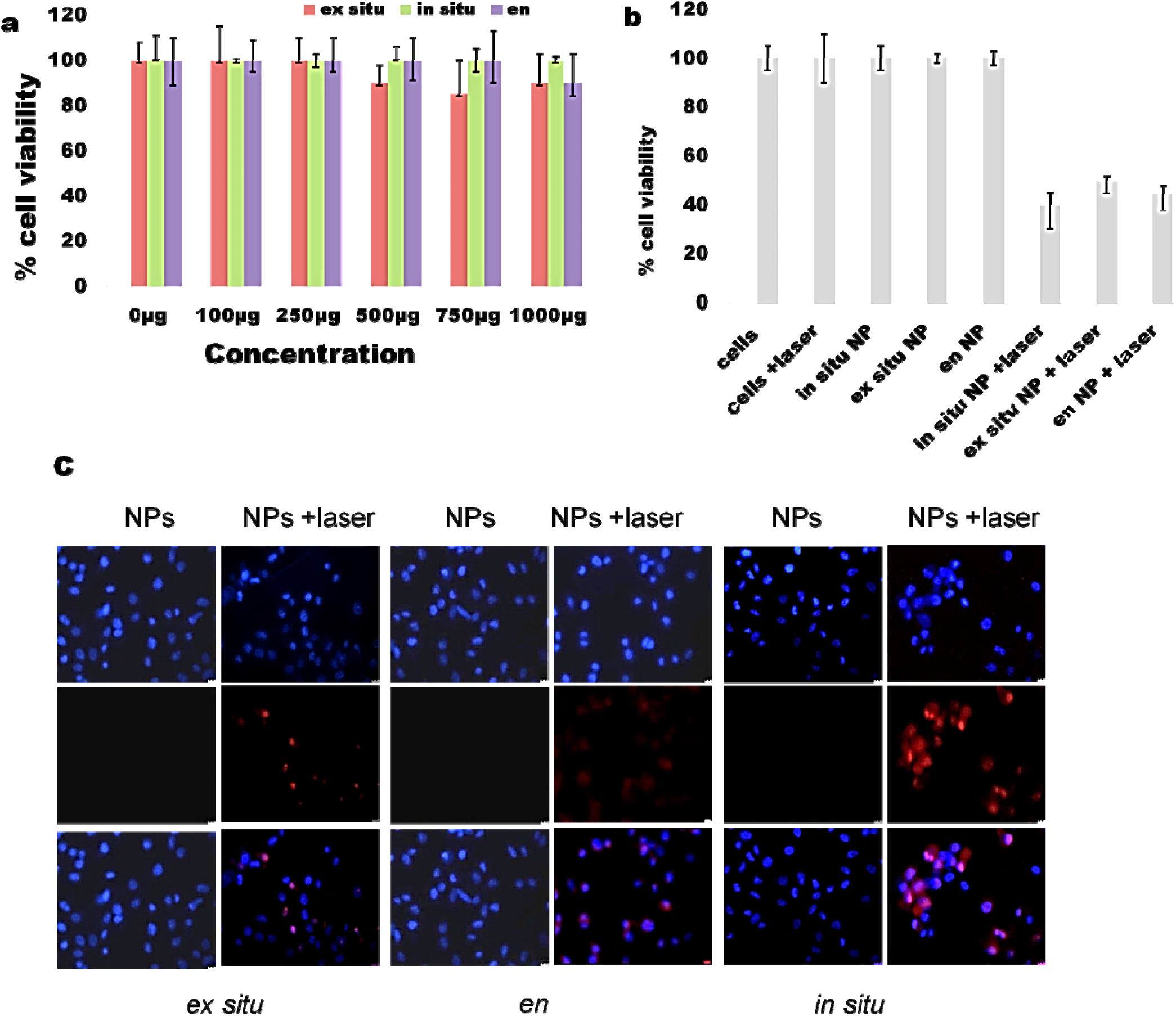
(a) Biocompatibility studies of FITC-PCLGC-IR nanoparticle in NIH 3T3 cells (b) Photothermal therapy of FITC-PCLGC-IR nanoparticle with 808nm laser in MDA-MB -231 cells (c) corresponding Hoechst 33342-Propidium Iodide staining of cell death in MDA-MB-231 cells before and after laser treatment.

### In vitro fluorescence stability of nanoformulation in MDA-MB-231 cells

The effect of laser on cell labeling with NPs was studied using confocal laser scanning microscope. All the three nanoformulation showed prominent labeling of MDA-MB 231 cells. Both *in situ* and *en* NPs showed good labeling in confocal imaging compared to *ex situ* NPs. Both *in situ* and *en* NPs nanoparticles are uniform in shape with high zeta potential which led them to interact with negatively charged cell surface for maximum uptake. The low cellular uptake of *ex situ* NPs can be attributed to aggregation of NPs upon FITC conjugation. A FITC conjugation can help in tracking and monitoring of the laser based treatment as IR 820 loaded NPs lose its imaging ability after photothermal therapy. Dyson et al have reported that FITC fluorescence increase upon hyperthermia in cells [25]. To visualize the effect of laser on FITC conjugation and its use as bioimaging molecules, we visualize the cells with nanoparticles before and after laser treatment (Fig. 7). The cells did not show any significant change in fluorescence before and after laser treatment. We also observed that hyperthermia did not affect the FITC fluorescence. All the formulation have the significant role in post laser imaging. An *ex situ* conjugation could be useful for on-demand delivery of imaging agent just before application Whereas, *in situ* conjugation and FITC encapsulation can be done during the synthesis of the nanoformulation to form stable NPs.

**Figure 7:**
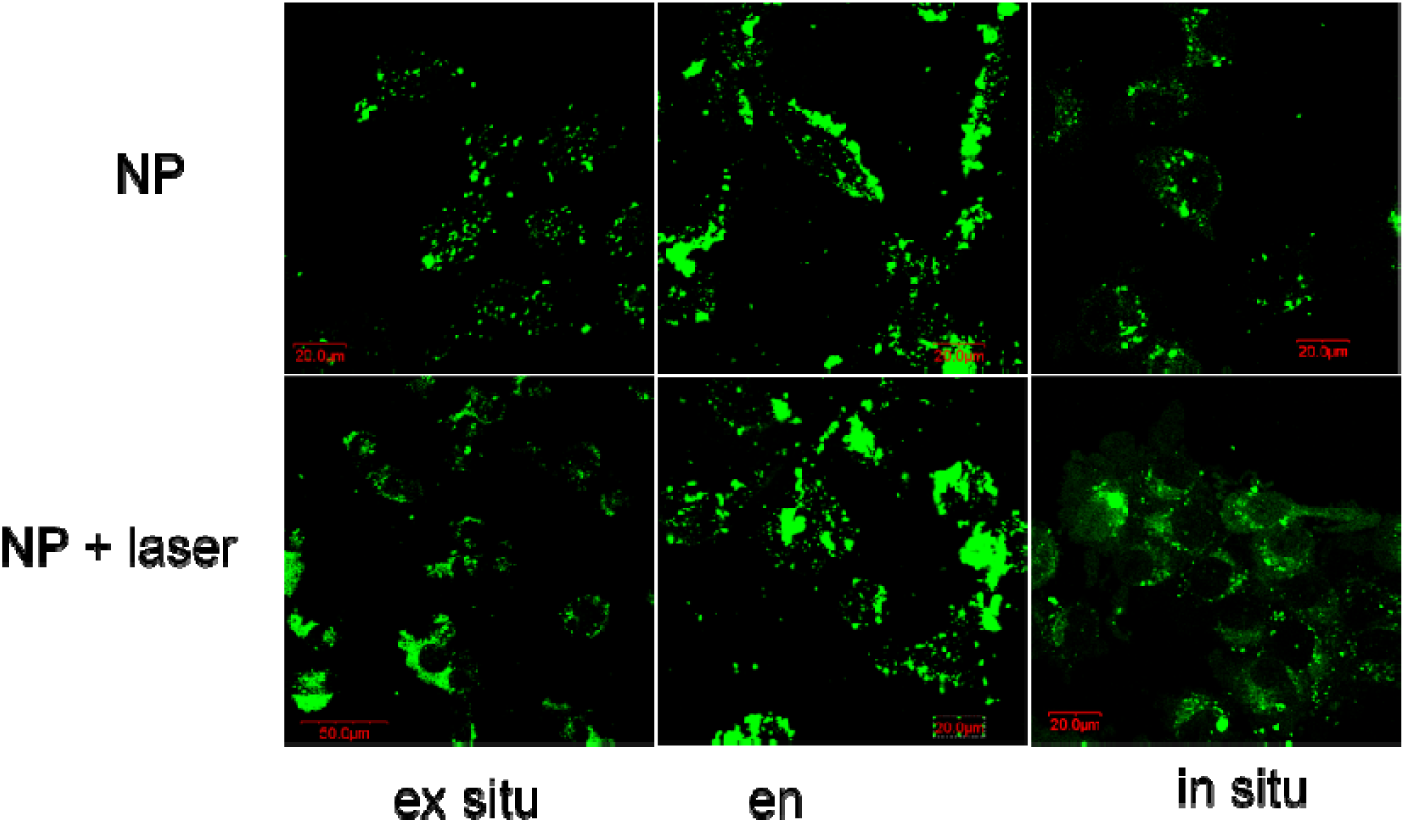
Confocal microscopic image of FITC-PCLGC-IR NPs before (upper) and after laser treatment (lower panel).

## Conclusions

In summary, we have performed the detailed study of in vitro stability of FITC conjugated NPs and its significant role in bioimaging and photothermal therapy. We have shown that how FITC conjugated PCLGC-IR NPs sorted out the limitation of theranostic dyes FITC fluorescence remains unaltered even after laser irradiation suggesting its role in bioimaging and in the vitro tracking of hyperthermia induced cell death. Also, FITC incorporation did not affect the photothermal property of IR 820 in nanoformulation for cancer therapy. A further study is required to check the similar effect with other dyes for monitoring the hyperthermia induced cell death in both in vitro and in vivo. In the nutshell, in vitro fluorescence stability of FITC conjugated NPs could be a promising approach for effective biocompatible and biodegradable image guided photothermal therapy.

## Acknowledgements

The authors would like to acknowledge SAIF-IITB and the chemical engineering department for characterization studies. P. K. is thankful to the Indian Council of Medical Research, New Delhi for a senior research fellowship [(File no. 3/ 1/3/JRF-2008/HRD-102(32238)]. The authors also acknowledge IITB-Healthcare for financial support.

